# The Colibactin-Producing *Escherichia coli* alters the tumor microenvironment to immunosuppressive lipid overload facilitating colorectal cancer progression and chemoresistance

**DOI:** 10.1101/2023.03.13.523827

**Authors:** Nilmara de Oliveira Alves Brito, Guillaume Dalmasso, Darja Nikitina, Amaury Vaysse, Richard Ruez, Lea Ledoux, Thierry Pedron, Emma Bergsten, Olivier Boulard, Lora Autier, Sofian Allam, Laurence Motreff, Pierre Sauvanet, Diane Letourneur, Gabriel Tang, Johan Gagnière, Denis Pezet, Catherine Godfraind, Michel Salzet, Emmanuel Lemichez, Mathilde Bonnet, Imène Najjar, Christophe Malabat, Marc Monot, Denis Mestivier, Nicolas Barnich, Isabelle Fournier, Sean P. Kennedy, Amel Mettouchi, Richard Bonnet, Iradj Sobhani, Mathias Chamaillard

## Abstract

Intratumoral bacteria locally contribute to cellular and molecular tumor heterogeneity that support cancer stemness through poorly understood mechanisms. This study aims to explore how Colibactin-producing *Escherichia coli* (CoPEC) flexibly alters the tumor microenvironment in right-sided colorectal cancer (CRC). Metabolomic and transcriptomic spatial profiling uncovered that CoPEC colonization establishes a high-glycerophospholipid microenvironment within the tumor that is conducive to exhaustion of infiltrated CD8^+^ T cell and has a lowered prognostic value in right-sided CRC. Mechanistically, the accumulation of lipid droplets in infected cancer cells relied on the production of colibactin as a measure to limit genotoxic stress and supply with sufficient energy for sustaining cell survival and lowering tumor immunogenicity. Specifically, a heightened phosphatidylcholine remodeling of CoPEC-infected cancer cells by the enzyme of the Land’s cycle coincided with a lowered accumulation of proapoptotic ceramide and lysophosphatidylcholine. Consequently, a reduced infiltration of CD8^+^ T lymphocytes that produce the cytotoxic cytokines IFN-γ was found where invading bacteria have been geolocated. By contrast, such an immunosuppressive dysmetabolic process was not observed when human colon cancer cells were infected with the mutant strain that did not produce colibactin (11G5δClbQ). This work revealed an unexpected property of CoPEC on lipid overload within tumors that could locally provide an inflammatory environment leading to immunosuppressive mechanisms and tumor expansion. This may pave the way for improving chemoresistance and subsequently outcome of CRC patients who are colonized by CoPEC.

## INTRODUCTION

Colorectal cancer (CRC) is the third most common form of malignancy and the second leading cause of cancer-related death worldwide.^1^ Patients with right-sided CRC have a worse prognosis than left-sided CRC and it has already been reported that they do not respond well to conventional chemotherapies.^2^ Chemotherapy drugs used to treat CRC primarily include oxaliplatin and cytotoxic drugs that inhibit the enzyme activity of thymidylate synthase. Whereas most patients with advanced CRC are primarily responsive to first-line chemotherapies, the 5-year survival rate is lower than 10% as a consequence of acquired chemoresistance. Only recently has been established that tumor-type features of the intratumoral microbiota are closely associated with resistance to chemotherapies and tumor recurrence.^3,4^ Specifically, some bacteria of often very low biomass may populate specific niches within the metastatic tumor microenvironment for impeding immune surveillance^5^ The use of patient-derived xenograft models revealed that such intratumoral microniches with *Fusobacterium nucleatum* are maintained even in distant metastases.^6^ This leads to the acquisition of chemoresistance through the modulation of autophagy. ^4^

Colibactin is a secondary metabolite produced by certain *Escherichia coli* strains that preferentially colonize the right-sided colon.^7,8^ Bonnet et al.^9^ reported a poorer prognosis outcome of patients that are colonized by Colibactin-producing *Escherichia coli* (CoPEC), which is detected in about 50 to 60% of human CRC biopsies compared to ∼20% of patients with diverticulosis. ^10,11^ This suggests that CoPEC may contribute to the recurrence of CRC. Accordingly, it has been established that the colibactin induces double-strand breaks (DSBs) ^12^ as well as generates DNA adducts ^13^ and genomic aberrations with an increased mutational load ^14,15^. This led to the identification of a specific DNA damage signature with an AT-rich hexameric sequence motif in tumors of patients that have been colonized by CoPEC strains^16^. Accordingly, colon tumorigenesis is accelerated in APC^Min/+^ mice that are colonized by CoPEC ^9,17^. Another work evidenced that organoids that recovered from short-term infection with CoPEC show characteristics of CRC cells including enhanced proliferation, Wnt-independence, and impaired differentiation that are often achieved by mutations in the gene encoding for Adenomatous polyposis coli (APC) ^18^. In APC^Min/+^ mice, colon tumorigenesis is accelerated following colonization by CoPEC through modulation of the autophagic pathway in intestinal epithelial cells ^17^. However, the potential effect of Colibactin on tumor recurrence and resistance to chemotherapy has not yet been examined in CRC.

Herein, we identified that colonization by CoPEC is an unfavorable prognostic factor in a subtype of poorly immunogenic right-sided colorectal tumors that are defined as consensus molecular subtype 3 with a lipid overload. Specifically, spatially-resolved metabolomics applied to tumors that are colonized by CoPEC revealed a significant intratumoral deposit of glycerophospholipids in areas that are populated by bacteria. Taken together, our findings clarify how Colibactin may establish tumor heterogeneity for evading immune surveillance. This provides unique insights for fostering the development of therapies targeting the oncogenic-driven lipid reprogramming that is induced by CoPEC.

## RESULTS

### Right-sided CRC tumors that are colonized by CoPEC are associated with cancer recurrence and a distinct microbiome composition

Microbiome profiling revealed that right-sided colon tumors are associated with a dense community of bacteria encased in a likely complex matrix that contacts the colon epithelial cells. ^19^ This led us to examine whether the composition of intratumoral microbiota could be linked to right-sided CRC recurrence that is mainly attributed to chemoresistance. To this end, we performed 16S rRNA gene sequencing on tumor samples of 76 patients with right-sided CRC from two independent cohorts. We first compared the respective influences of a denoising method and a clustering method on our 16S rRNA gene amplicon dataset. Given that no discrepancy was obtained on the most abundant bacteria with both methods (data not shown), we investigated the biodiversity of a total 3,511 amplicon sequence variants (ASVs) that were produced by the DADA2 method after filtering. PCoA projection of the data revealed compositional differences between tumoral tissues from patients with relapse when compared to those without recurrence (Figure 1A and 1B). Adonis analysis showed significant differences between the two study groups both at genus and species taxonomic level (p value= 0.0066, F model = 2.35 and p value= 0.0034, F model = 2.23, respectively) (Figure 1C and 1D). Among the differentially abundant genera, we found that the abundance of bacteria-related genus *Fusobacterium* was significantly more prevalent in tumors from patients in remission. By contrast, a large subset of recurrent tumors harbored predominant populations of bacteria that belong to the genus *Escherichia-Shigella*. Accordingly, the specie *Escherichia coli* was the most enriched in recurrent CRC tissues as compared to those from patients without relapse (Figure 1D). Based on these differences, we further investigated whether the presence of *pks* within the tumor could influence survival outcomes, particularly in patients with stage III and stage IV. The detection of the genomic island *pks* was validated by two different PCR-based methods on DNA that was extracted from right-sided CRC tissues. This revealed that right-sided CRC patients colonized by CoPEC are associated with poor survival (p=0.018) (Figure 1E). This is in agreement with the worst prognosis that has been documented in patients who are colonized by CoPEC.^9^ Given that the genotoxin colibactin shapes gut microbiota in mice,^20^ we sought to assess whether the heterogeneity of the intratumoral microbiota is statistically distinct between CoPEC-positive and -negative tumors. Interestingly, a significantly lowered Shannon’s alpha diversity index was observed in tumors that were not colonized by CoPEC (Figure 1F). Likewise, CoPEC colonization significantly lowered the bacterial richness in the ASV-based dataset as quantified by the Simpson index (Figure 1F). This was in agreement with other alpha diversity estimates even though it failed to reach significance (Supplementary Figure S1). These results suggested us that CoPEC colonization impacts both the evenness and richness of the composition of right-sided CRC tumors. This led us to evaluate the influence of CoPEC colonization on patterns in beta-diversity. Interestingly, principal component analysis of beta-diversity revealed 35 bacterial taxa statistically distinct between the groups (Figure 1G). Unlike what is observed in CRC liver metastases that are mainly populated with *Fusobacterium nucleatum*,^5^ only the taxa *Escherichia-Shigella* was significantly more abundant in the right-sided tumors from patients colonized by CoPEC. Along with the *Escherichia-Shigella* group, nine bacterial ASVs linked to *Prevotella*, *Bacteroides* (*B. uniformis, B. fragilis, B. stercoris* and *B. salyersiae*), *Lachnoclostridium, Barnesiella,* and *Odoribacter splanchnicus* were significantly increased in the CoPEC positive group. At the same time, we detected 15 ASVs that were significantly depleted in the group of CoPEC-positive tumors (Figure 1G). Among those that may provoke or participate in inflammatory processes or carcinogenesis, the abundance of *Acidaminococcus* was also decreased in CRC patients’ stool samples compared to colorectal adenoma and control group.^21^ The analysis of consensus molecular subtypes (CMS) based on bulk RNA-seq analysis revealed that more than 50% of tumors colonized by CoPEC were classified as CMS3 that have metabolic dysregulation with higher activity in glutaminolysis and lipogenesis. Next, we selected only patients from the CMS3 group and performed differential abundance analysis according to their CoPEC status (Figure Supplementary Figure S2). Similar to previous results, we detected an enrichment of six bacterial taxa that were previously associated with development of intestinal inflammatory processes. Specifically, *B. uniformis*, *O. splanchnicus, Bacteroides plebeius, Desulfovibrio piger* and *Veillonella* were found more abundant in the CoPEC-positive tumors. Among those, *B. plebeius* was over-represented in Lynch syndrome patients^22,23^ and *Veillonella* were found to be presented only in CRC patients. ^24^ These results could indicate that the processes of inflammation induced by intratumoral bacteria can take place in different ways in response to CoPEC colonization

**Figure 1.**
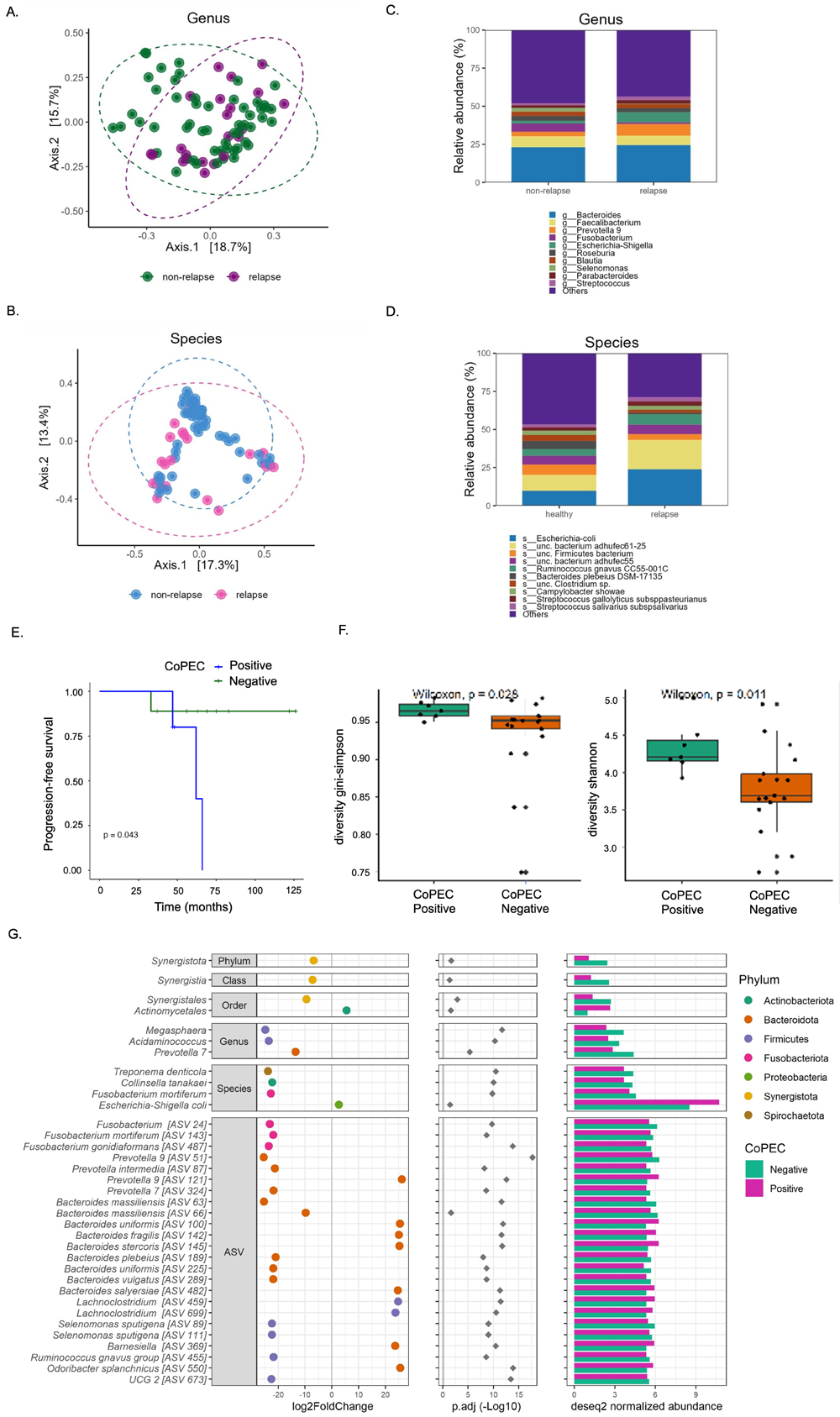
Right-sided CRC patients differ in relation to bacteria composition and showed poor survival in patients colonized by Colibactin. Principal Coordinate Analysis (PCoA) with Bray-Curtis distances matrix based on Deseq2 normalized data between non-relapse and relapse right-sided CRC patients in relation to genus (A) and species (B). The relative abundance between non-relapse and relapse right-sided CRC patients in relation to genus (C) and species (D). (E) Kaplan-Meier analysis showed that right-sided CRC patients colonized by Colibactin-producing *Escherichia coli* (CoPEC)-positive predict poor survival (patients with stages I and II were excluded from this analysis). (F) Alpha-diversity analysis between positive and negative CoPEC patients. (G) Beta-diversity analysis results between positive and negative CoPEC patients. The left plot represents the log_2_ fold change value of the mean normalized abundance ratio of positive CoPEC vs. negative CoPEC; the middle plot represents the adjusted p-value obtained by Deseq2 differential abundance analysis; the right plot represents the mean normalized abundance for each significant abundance bacteria.

### Enrichment of transcripts linked to lipid metabolism in relapsing tumors from patients colonized by CoPEC

The greater abundance of bacteria related to *Escherichia-Shigella* in relapse patients led us to explore the transcriptomic landscape in the different groups in relation to CoPEC colonization from patients with relapse as well as those without recurrence (non-relapse group). Differential gene expression analysis between CoPEC-positive and -negative tumors from patients with relapse revealed 547 up- and 869 down-regulated genes. On the other hand, CoPEC-positive and -negative tumors from patients without recurrence showed 151 up- and 136 down-regulated genes (Figure 2A). By performing functional mapping and annotation (FUMA), biological processes gene ontology (GO) term analysis was presented in Supplementary Figure S3, among which cell division, cell cycle process, cellular response to stress, phospholipid and glycerophospholipid process, DNA damage stimulus and others were closely associated with Colibactin in CRC progression. For instance, up-regulated genes in CoPEC-positive tumors from the relapse group were enriched in the cellular lipid metabolic process and double-strand break repair (Figure 2B and 2C). Especially, mRNA levels associated with the regulation of lipid metabolism were overrepresented. Among them, diacylglycerol kinase gamma (DGKG), a gene that encodes an enzyme that generates phosphatidic acid (PA) by catalyzing the phosphorylation of diacylglycerol (DAG) was up- regulated in tumors colonized by CoPEC in the relapse group. We also observed that Lysophosphatidylcholine acyltransferase 2 (LPCAT2), which plays a role in phospholipid metabolism in the conversion of lysophosphatidylcholine (LPC) to phosphatidylcholine (PC) in the presence of acyl-CoA was increased in these same patients. In addition, different changes in the phospholipase A2 enzyme family genes, that catalyze the hydrolysis of the sn-2 position of membrane glycerophospholipids generating free fatty acids and lysophospholipids were identified. For example, PLA2G16, PLA2G4D and PLA2G2F were significantly up-regulated in tumors populated by CoPEC. In contrast, the transcript level of *PLA2G6* was down-regulated in this group. Interestingly, these alterations were accompanied by overexpression of transcripts implicated in the ceramide metabolism such as alkaline ceramidase 2 (ACER2) and sphingomyelin synthase 1 (SGMS1). The ceramidase ACER2 hydrolyzes long-chain ceramides to generate sphingosine while SGMS1 catalyzes in the forward reaction of transferring the phosphocholine head group of PC onto ceramide to form sphingomyelin (Figure 2D). Together, these lipid enzyme alterations encouraged us to investigate if Colibactin-producing bacteria may be involved in the lipid metabolic reprogramming of recurrent right-sided tumors.

**Figure 2.**
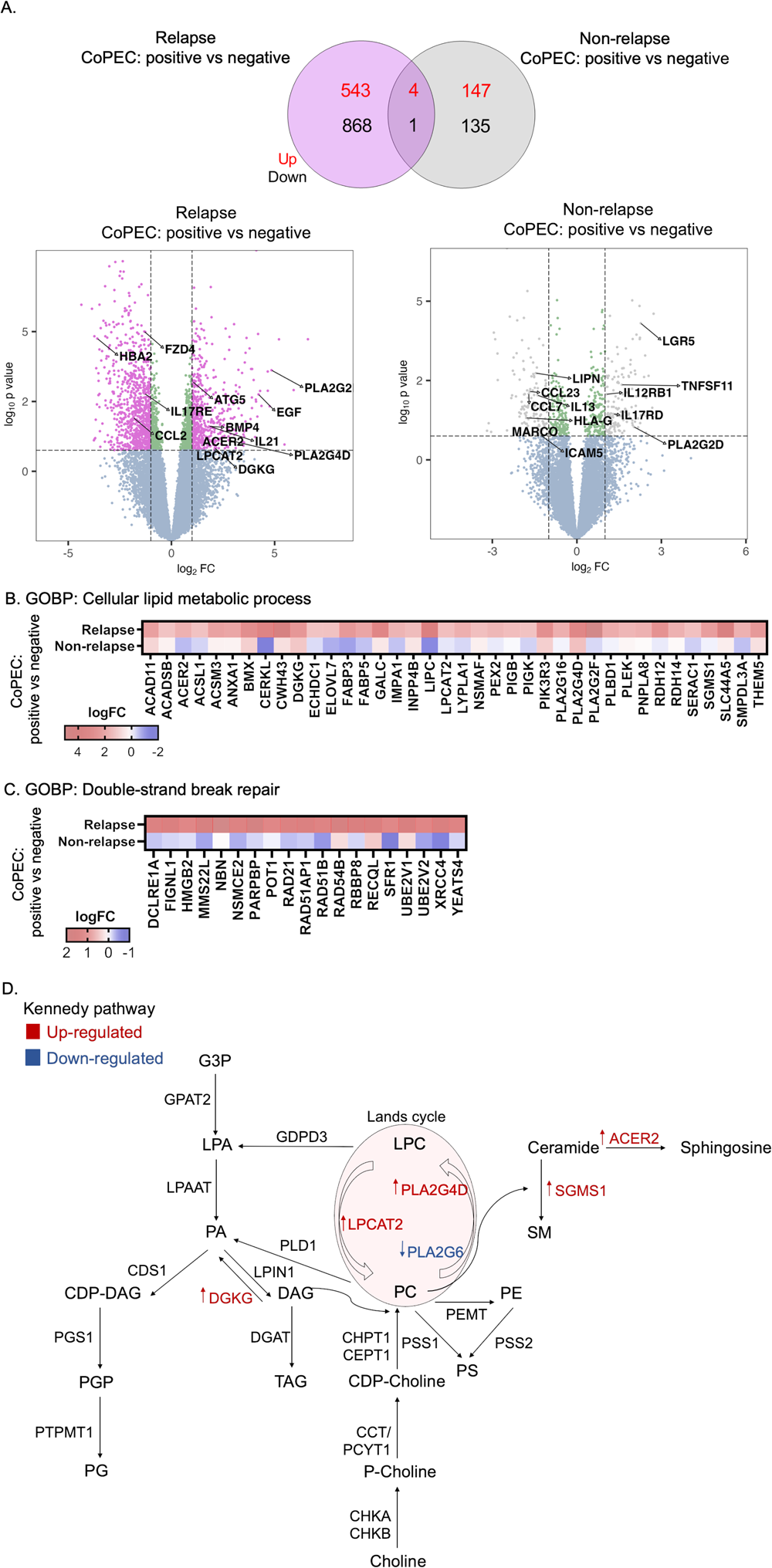
Patients colonized by CoPEC with relapse reveal enrichment of genes associated with lipid metabolism and DNA double-strand break. (A) Volcano plots and Venn diagram of differentially expressed genes in response to CoPEC-positive versus - negative tumors from relapse (n=17) and non-relapse groups (n=30). Dashed lines indicated the following significance threshold: 1.0 > log_2_ fold change < -1.0 and p-value < 0.05. (B) and (C) Heatmap showing logFC of genes associated with cellular lipid metabolic process and double-strand break repair (FUMA-GO biological process) in response to CoPEC- positive versus -negative tumors from relapse and non-relapse groups. (C) Schema showing genes involved in the biosynthesis of the lipid in both the Kennedy pathway (De novo synthesis) and Lands cycle. Highlighted genes in response to CoPEC-positive versus - negative tumors from relapse group. Red: up-regulated and Blue: down-regulated.

### Spatially-resolved metabolomic approach unveils bacterial regions that are highly metabolically active in response to CoPEC colonization

The response of gene expression associated with lipid metabolism in CoPEC-infected tumors led us to study the potential consequences of metabolic reprogramming of areas that are invaded by bacteria. To this end, spatially resolved metabolomics was applied on the 12 right-sided CRC tissue samples using a high spectral resolution of the 7T-MALDI-FTICR. By using *in situ* hybridization (FISH) imaging, we visually confirmed the heterogeneous geolocation of bacterial microniches in right-sided colorectal tissues. On each tissue, 3–13 specific regions of interest (ROI) were selected according to the FISH staining (Figure 3A). This resulted in a total of 90 ROI with bacteria microniches and 86 ROI without bacteria. Here, we detected 12 up-regulated metabolites in ROI with bacteria compared to ROI without bacteria within CoPEC-positive tumors (Figure 3B). Moreover, comparisons between ROI with and without bacteria in CoPEC-undetected tumors revealed 20 metabolites down and 11 over-expressed (Figure 3C). Specifically, 11 metabolites were found up-regulated only in patients colonized by CoPEC such as Benzenoids (n=1), Fatty Acyls (n=1), Glycerophospholipids (n=3), Organic acids (n=1), Organoheterocyclic (n=2), Sphingolipids (n=1) and Sterol Lipids (n=2)). In addition, metabolomic differential expression in ROI with bacteria microniches revealed a predominance of lipids (approximately 40%) in the CoPEC- positive compared with the CoPEC-negative right-sided tumors. Among those lipids identified, approximately 82% belong to the glycerophospholipids subclass such as phosphatidic acid (PA), phosphatidylcholine (PC), phosphatidylinositol (PI), phosphatidylethanolamine (PE), phosphatidylserine (PS), phosphatidylglycerol (PG) and others (Figure 3D). Subsequently, we performed a comparative analysis of metabolite intensity for the same m/z in all tumor tissue and the areas with bacteria. This showed that most of the metabolic heterogeneity is explained by microniches populated with bacteria (Figure 3E). Given that disturbances in the gut microbial profiles/communities are associated with heightened *de novo* lipogenesis, this led us to suggest that CoPEC may be responsible for metabolic dysregulation that promotes carcinogenesis.

**Figure 3.**
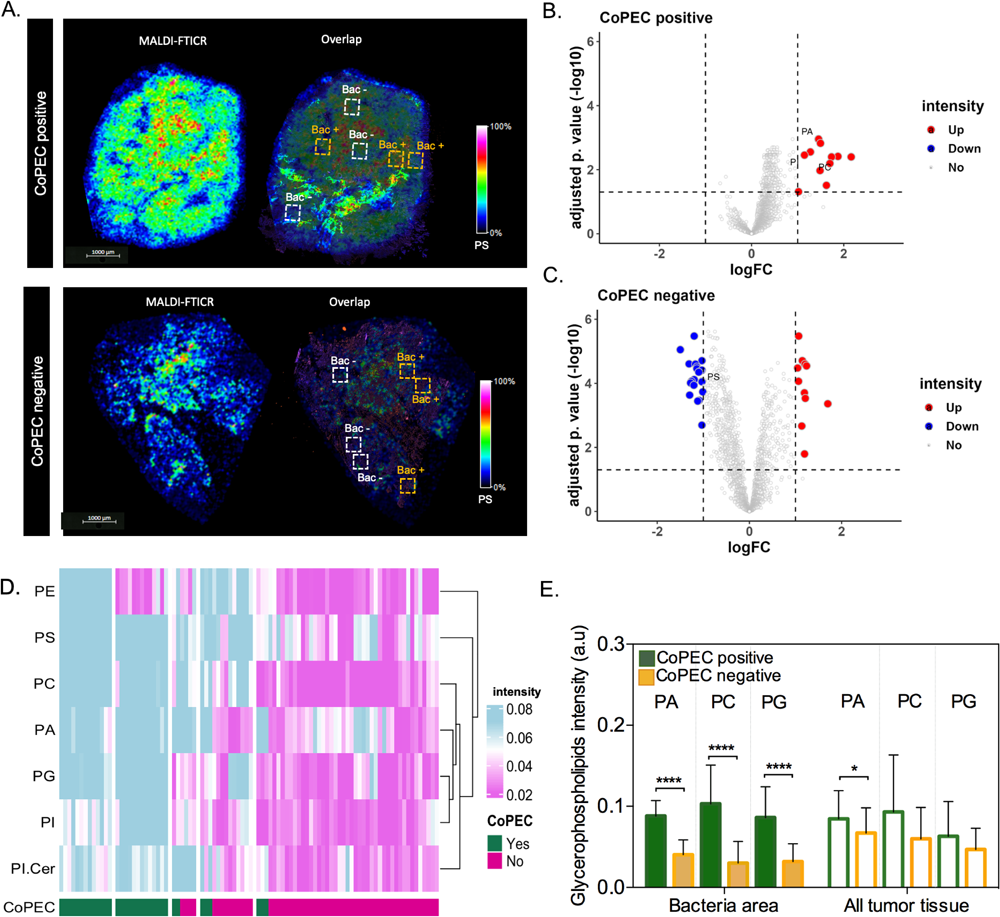
Spatial metabolomics revealed an increase in glycerophospholipid in tumor-associated bacterial microniches mainly in patients colonized by Colibactin-producing *Escherichia coli* (CoPEC). (A) Representative images of MALDI-FTICR and overlap with FISH (bacteria stained with general rRNA probe EUB338 conjugated to Alexa 555 - orange) from CoPEC-positive and -negative RCC tumors. (B) and (C) Volcano plot displaying metabolomic analysis from CoPEC-positive (ROI with bacteria versus ROI without bacteria) and negative CoPEC (ROI with bacteria versus ROI without bacteria) patients, respectively. Root means square normalization was applied, adjusted p. value (FDR correction) < 0.05, and log_2_FC > 1 or < -1 were considered. (D) Heatmap showing differentially expressed lipids in the ROI with bacteria between CoPEC-positive and -negative tumors. (E) Graphic showing the difference in metabolite expression in bacteria area (n=12) and in all tumor tissue (n=65).

### Lipid droplet accumulates in colon cancer cells that are infected by CoPEC

To assess whether colibactin may directly trigger lipid overload, human colon carcinoma cells HCT116 were infected with CoPEC strain (11G5) isolated from a patient CRC or a mutant strain that does not produce Colibactin (11G5Δ*clbQ*).^25^ In agreement with our spatial metabolomic analysis, CoPEC infection leads to lipid droplet accumulation and an increase of LPCAT2 levels (Figure 4A-4C). SpiderMass technology was then applied to define which glycerophospholipids contribute to the discrimination between 11G5-, 11G5ΔclbQ-infected and non-infected HCT116 cells. This led us to identify several differentially abundant glycerophospholipids such as PC, PS, PE and PI and decrease of LPC in response to CoPEC infection (Figure 4D-4H). By contrast to what was observed in response to cellular stress by chemotherapy, the level of ceramide was significantly reduced in response to CoPEC (Supplementary Table S1). Similar results were obtained with the syngeneic colorectal carcinoma cell line MC38 (Supplementary Table S2). Together, the CoPEC-induced lipid droplet accumulation was preceded by a heightened *de novo* lipogenesis, which encouraged us to continue investigating the mechanisms of how tumor-associated CoPEC can promote lipid overload.

**Figure 4.**
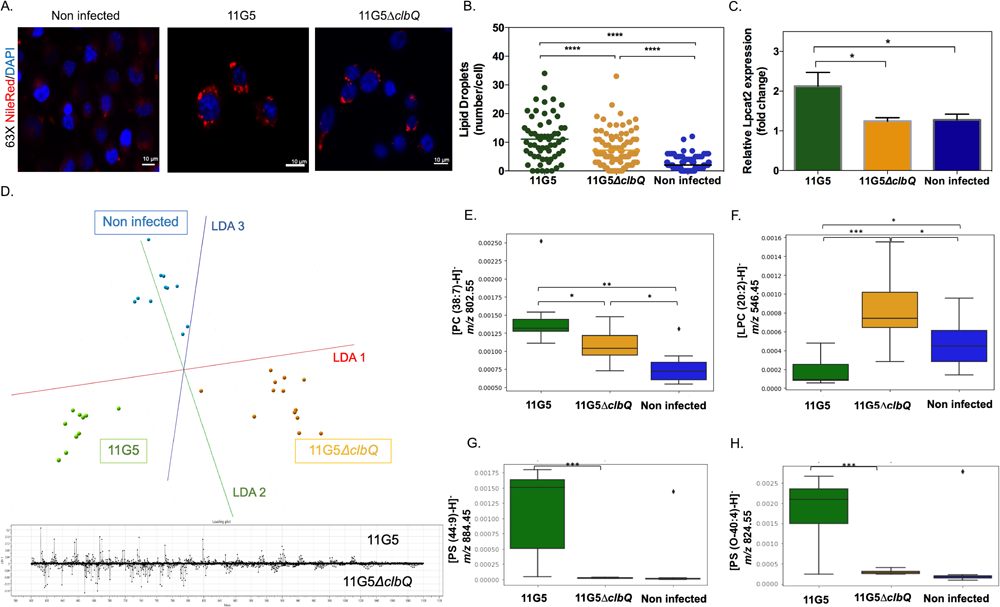
Bacterial Colibactin leads to lipid droplet accumulation in human colon cancer cells. (A) Representative image of Nile red staining (×63 magnification, scale bar[=[10[µm) using HCT116 cells. Nuclei (DAPI -blue), lipid droplet (red). (B) Lipid droplet quantification (300 cells) under different bacterial infections: CoPEC 11G5 strain, isogenic mutant 11G5Δ*clbQ* strain and negative control (NC, non-infected). (C) MC38 cells were infected with the 11G5 strain or the 111G5Δ*clbQ* and were analyzed 5 days post-infection. Lpcat2 mRNA levels from MC38 cells were quantified using qRT-PCR. (D) The built PCA-LDA classification model is based on three groups: HCT116 cells infected with CoPEC strain (11G5), a mutant strain that does not produce CoPEC (11G5ΔclbQ) and non-infected (negative control). Representative m/z chromatograms. (E - H) Graphs with the intensity of PC, LPC and PS differentially expressed under the conditions studied.

### CoPEC-induced lipid droplet accumulation and chemoresistance are preceded by elevated levels of reactive oxygen species

RNA-seq analysis was then performed to further characterize the early stress response of cancer cells to CoPEC that precedes lipid droplet accumulation. Applying the edgeR method revealed a total of 98 genes that were differentially expressed in either direction between cells that were infected by either CoPEC or its mutant that is unable to produce colibactin (p<0.0001). Among those, 94 were significantly up-regulated as displayed with a Volcano plot (Figure 5A). Over-representation analysis revealed that several differentially expressed genes were functionally related to processes that are related to both oxidative phosphorylation and adipogenesis, including Cyc1 which encodes a subunit of the cytochrome bc1 complex and Suclg1 that encodes Succinate-CoA Ligase GDP/ADP-Forming Subunit Alpha (Figure 5B-5D). These results suggested us that CoPEC may locally promote the accumulation of lipid droplets through induction of hypoxia and extracellular acidosis. Since lipid droplet production can act as switches in response to imbalances in energy metabolism and redox homeostasis ^27^, we next investigated the earlier accumulation of reactive oxygen species (ROS) in both MC38 and HCT116 cells. As shown in Figure 5E-G, a semi-quantitative assessment of the percentage of cells with CellRox was performed (score 0 to 3, see Material and Methods section) and a notable increase in cells with high fluorescence (score 3) was observed after CoPEC infection. Given that elevated ROS may result in immunogenic cell death, we hypothesized that the accumulation of lipid droplets may be a protective mechanism that curtails the efficacy of anticancer drugs. This led us to evaluate whether CoPEC-induced ROS formation could affect the sensitivity of CRC cells to oxaliplatin which is a third-generation diaminocyclohexane-containing platinum compound. Accordingly, CoPEC infection remarkably lowered the IC50 of oxaliplatin in HCT116 cells (Figure 5H). This is in agreement with previous studies showing that the level of enzyme supporting PC synthesis is enhanced in oxaliplatin-resistant cells compared to untreated parental cells.^1,2^

**Figure 5.**
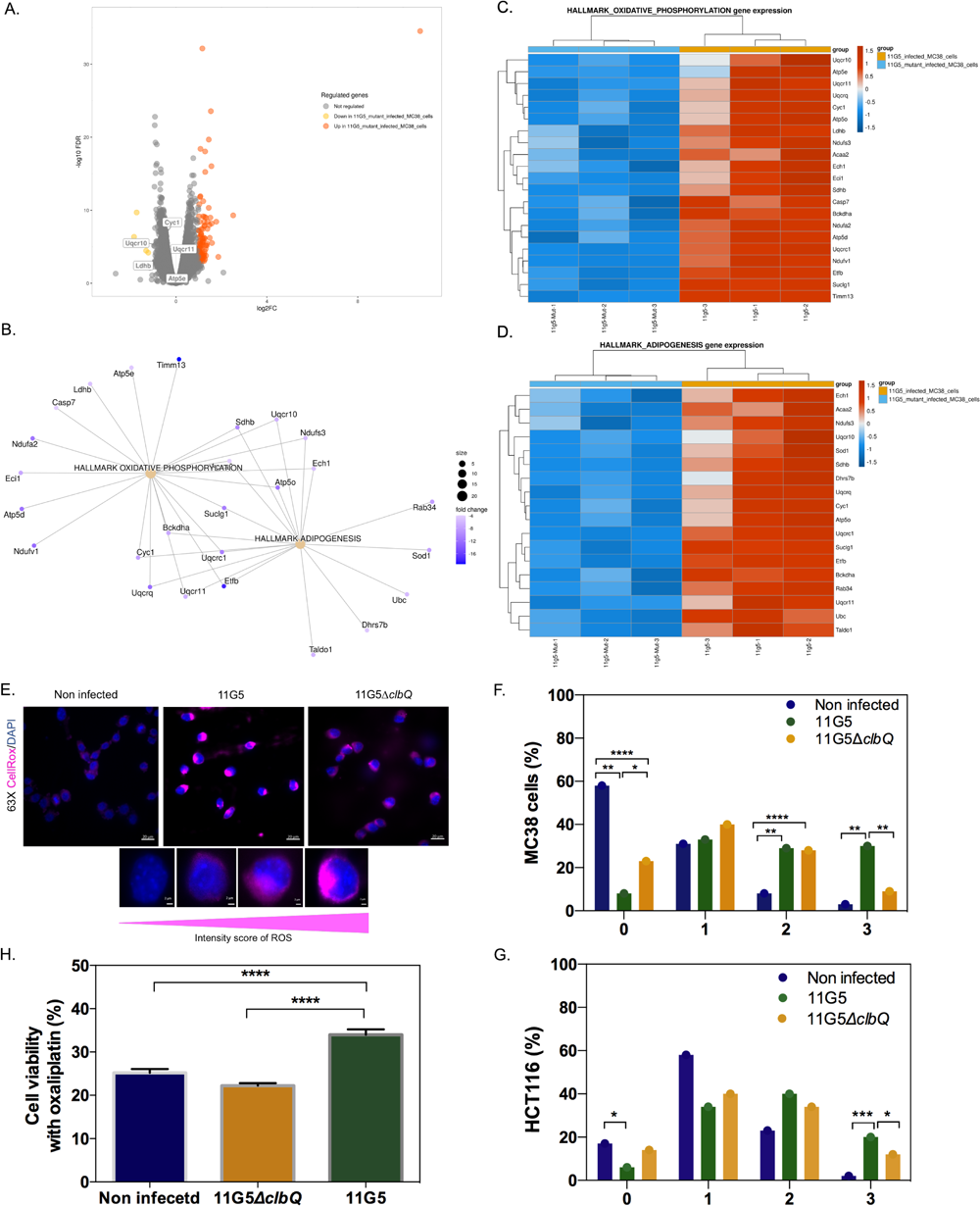
Chemoresistance in CRC cells is preceded by elevated levels of reactive oxygen species after CoPEC colonization. (A) Volcano plot representation of differentially expressed genes between MC38 infected with a mutant strain that does not produce CoPEC (11G5ΔclbQ) and CoPEC strain (11G5). (B-D) Representation of the enrichment of genes involved in both oxidative phosphorylation and adipogenesis. (E) Representative image of CellRox staining for intracellular reactive oxygen species in HCT116 infected with CoPEC strain (11G5), a mutant strain that does not produce CoPEC (11G5ΔclbQ) and non-infected (negative control). Fluorescent images were captured after 4h of treatment (3h of infection + 1h pos-infection) under MOI 10. (F and G) A semi-quantitative assessment of the percentage of ROS in 300 cells was evaluated using different scores from 0 to 3 using MC38 and HCT116 cells, respectively. Data are expressed by the percentage of cells in each score and group (two independent experiments and duplicates for each experiment). * p < 0.05, ** p < 0.01, *** p < 0.001 and *** p<0.0001. (H) HCT116 cells were infected with CoPEC strain (11G5), a mutant strain that does not produce CoPEC (11G5Δ*clbQ*) and non-infected supplemented with oxaliplatin (5 μg/mL). Cell viability was assessed using WST-1 Assay Reagent - cell proliferation assay. Non-infected cells without oxaliplatin were used to represent 100% viability. Values represent means ± SEM.

### CoPEC-induced lipid droplet accumulation impairs the immunogenicity of right-sided CRC

This led us to the hypothesis that the formation of lipid droplets may lower immunogenicity by supporting acidosis-driven epithelial-to-mesenchymal transition. Accordingly, bulk RNAseq analysis from tumor tissues unveiled a down-regulation of several genes that are related to B cell activation in CoPEC-positive tumors, such as CD19, CCR6, CD40LG and DOCK11 (Figure 6A). Accordingly, a significant enrichment of cytotoxic CD8^+^ T-cells and B-cells proportion was observed in CoPEC-positive tumors when using the web-accessible TIMER2.0 algorithm (Figure 6B). This led us to determine whether there is a difference in CD8^+^ T-cell infiltrates in intra- and inter-tumor annotated as regions of interest (ROI) with or without bacteria. Accordingly, RNAscope-FISH overlay images revealed that bacteria-populating microniches are characterized by immunosuppressive effect with a decrease of tumor-infiltrating CD8^+^ T-cells, mainly in patients colonized by CoPEC. In addition, in both groups, we detected a significant reduction of CD8^+^ T-cells in ROI with bacteria when compared to ROI without bacteria (Figure 6C-6E). Analyzing specific CD8^+^ T-cells that produce IFNγ, we observed that this tendency is sustained among tumors that are colonized by CoPEC, reinforcing that the effect of the tumor-associated bacteria is localized (Figure 6F). By applying spatial profiling approaches, we were able to quantify the areas of overlap between CD8^+^ T-cells and glycerophospholipids. Interestingly, we observed a significant inverse correlation between the quantification of CD8^+^ spots and PC intensity (r= - 0.56, p= 0.04) in the CoPEC patients (Figure 6G). By contrast, this correlation was lost in CoPEC-negative tumors (r=-0.04, p=0.81) (Figure 6H), suggesting CD8^+^ T-cell expression is tightly intertwined with a specific metabolic pathway in a Colibactin-induced immune-suppressive microenvironment in right-sided CRC. Accordingly, a negative correlation between IFNγ was observed in tumors that were subcutaneously implanted in wild-type mice that were colonized by CoPEC (Figure 6I).

**Figure 6.**
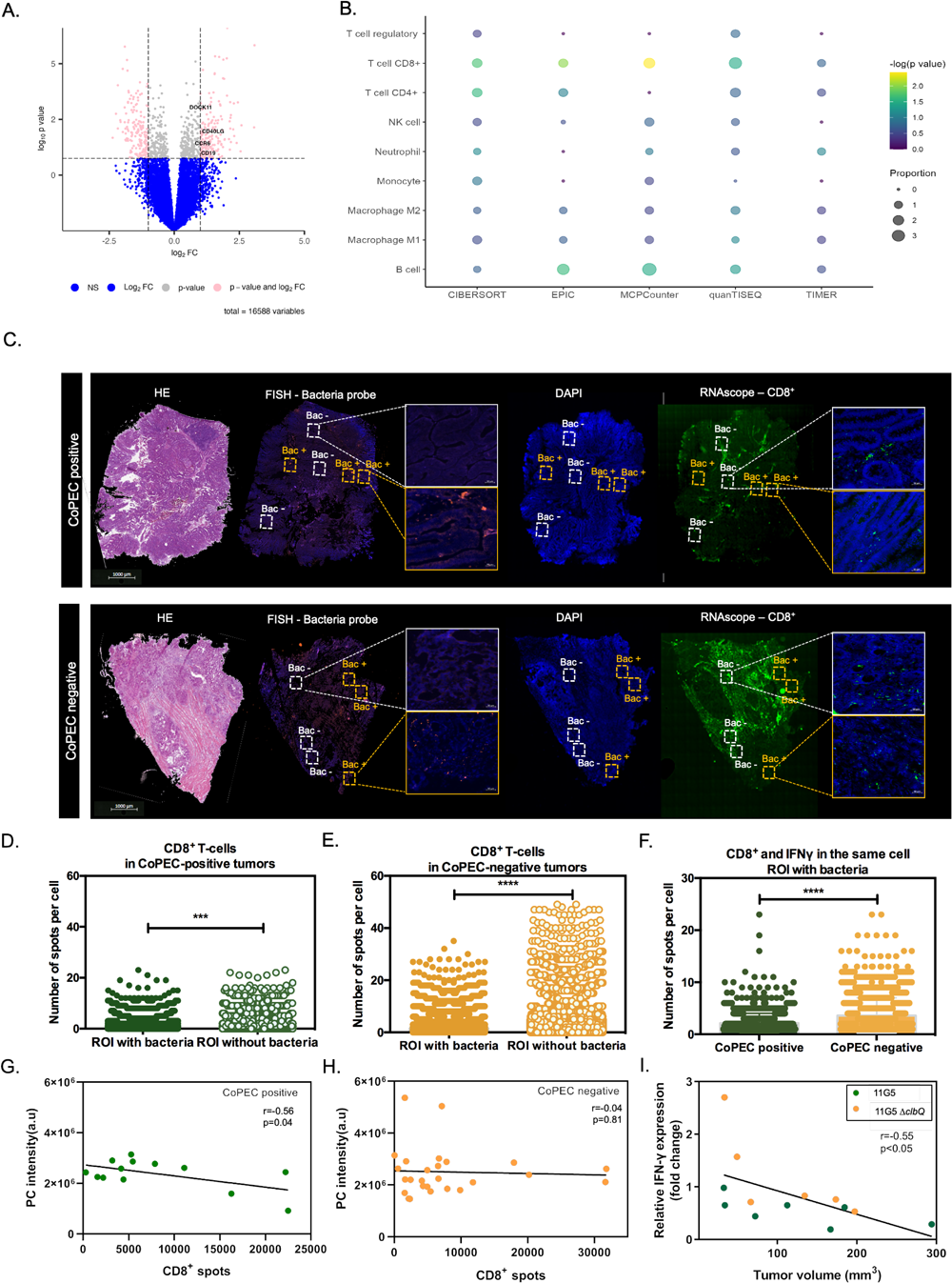
Bacterial microniches are poorly infiltrated with IFNγ-producing CD8^+^ T-cells in right-sided CRC patients colonized by CoPEC. (A) Volcano plot representation of differentially expressed genes between CoPEC-positive and -negative patients. Dashed lines indicated the following significance threshold: 1.0 > log_2_ fold change < -1.0 and p-value < 0.05. (B) Bubble plot showing the proportional difference of immune cells between CoPEC- positive and -negative patients based on the following computational methods: CIBERSORT, EPIC, MCP-counter, quanTIseq and TIMER. (C) Representative images of H&E, *in situ* hybridization (FISH - bacteria stained with general rRNA probe EUB338 conjugated to Alexa 555 - orange) and the nuclear DNA stained with DAPI (blue)), RNAscope (Hs-CD8A (560391-C3, Opal 520 - green) from CoPEC-positive and -negative RCC tumors. White square: ROI without bacteria and orange square: ROI with bacteria. (D) and (E) Quantification of CD8^+^ T-cells in the ROI with and without bacteria CoPEC-positive and - negative RCC tumors, respectively. (F) Quantification of CD8^+^ T-cells and IFNγ in the same cells in the ROI with bacteria from CoPEC-positive and -negative tumors. *** p < 0.001 and **** p<0.0001. (G) and (H) Pearson correlation between CD8^+^ T-cell spots detected by the RNAscope and phosphatidylcholine (PC) identified by the MALDI-FTICR in CoPEC- positive and negative RCC patients, respectively. (I) Pearson correlation between IFNγ and tumor volume from animals infected with 11G5 strain (n=7) or 11G5ΔclbQ strain (n=6).

## DISCUSSION

It remains a prerequisite to defining how inter-tumor heterogeneity and differences of the intra-tumoral microbiome can lead to variable responses to chemotherapies. A majority of detected bacteria (6 out of 10) in this group were previously associated with the development of intestinal inflammation, CRC, or other inflammatory bowel disease (IBD). For instance, *Bacteroides fragilis* has *B. fragilis* toxin (bft) gene that produces fragilysin or BFT. BFT disrupts tight contacts, and increases the permeability and damage of the intestine, thus being involved in the development of inflammation and carcinogenic processes.^28,29^ Another *Bacteroides* species B*. uniformis* and *B. stercoris* previously were found to be significantly enriched in participants with bloody stools and were associated with higher risk to develop CRC.^30^ *Lachnoclostridium* species have been presented as promising bacterial markers for the non-invasive stool-based diagnosis of colorectal adenoma.^31^ *Barnesiella* species were found to be associated with systemic inflammation.^32^ *Odoribacter* species in the previous studies have been correlated with somatic mutations and cell proliferation in CRC patients alongside decreased abundance in cystic fibrosis, non-alcoholic fatty liver disease, and IBD.^33–35^ Next, we selected only patients from the CMS3 group and performed differential abundance analysis according to their CoPEC status. In this group, we also detected a reduction in the number of commensal bacteria including *Lachnospira*, *Ruminococcus bromii*, *Anaerostipes*. *Lachnospir*a is considered as anti-inflammatory and healthy colonocytes promoting bacteria;^36^ *R. bromii* is a common member of the human gut and can degrade resistant starches that are present in human diets thus energetically supporting other commensal bacteria.^37^ *Anaerostipes* species are among the most active lactate consumers in the human colon, they also are short chain fatty acids producers.^38^ Compared with the results obtained from the general cohort, given results on CMS3 patients’ group shows greater decrease in number of commensal bacteria with established anti-inflammatory properties. These results suggest that the effect of colibactin on CMS3 is exacerbated and leads to a decrease in bacterial diversity normally associated with benefits for human health.

Previous works have shown that CoPEC induces double-strand DNA breaks, DNA mutations and can modulate the tumor microenvironment to favor the emergence of senescent cells, affecting tumor progression. ^8,12,25^ In this context, bulk analysis obscured our understanding of how host-microbiota is locally regulated in solid tumors because this approach cannot provide insights into the molecular events that locally occurred either sequentially or in parallel throughout the proliferation of resistant tumor cells ^5,39^. This paradigm led us to apply state-of-the-art high-throughput *in situ* spatial profiling technologies for defining whether CoPEC may locally contribute to metabolic heterogeneity and consequently may create a potential vulnerability to anti-cancer drugs. Our study identified that heterogeneous bacteria spatial distribution including *Escherichia coli* affects lipid metabolism in the tumor microenvironment. Specifically, the intra-tumoral areas that are populated by Colibactin-producing bacteria contribute to unbalanced lipid metabolism that may provide the breeding ground for the emergence of resistant cancer cells. Accordingly, the transcript level of key enzymes of the Land’s cycle that was particularly correlated with PC, PA, and ceramide metabolism. Besides, we identified differences in PA, PC, PI, PE, PS, and PG intensity in bacteria microniches between CoPEC-positive and -negative groups. Of note, PC and PI are generally increased in CRC and have been associated with cancer development and progression ^46,47^. Furthermore, PS can be synthesized from PC and PE and plays an important role in mitochondrial function and apoptosis ^48^. In addition, glycerophospholipids also play an important role in the autophagy pathway ^49^. Accordingly, lipogenesis is heightened in cancer cells that were infected by CoPEC but not by mutant bacteria that do not produce colibactin. Lipid droplet serve as an energy reserve for cancer cells that require a substantial amount of energy for their rapid growth and proliferation. Of note, a recent work described that *Fusobacterium nucleatum* promotes CRC cancer cells to acquire stem cell[like features by lipid droplet[mediated Numb degradation. ^50^ In our study, Spider Mass technology revealed that PC is a main structural component of lipid droplets in CoPEC HCT116 cells. The CoPEC-induced dysregulation in lipid metabolism may affect a number of cell functions. Our hypothesis that lipid alteration may lower immunogenicity was clarified applying *in situ* spatial analysis that demonstrated a decrease of lowered infiltration of IFNγ-producing CD8^+^ T-cells in local bacteria microniches, mainly in patients colonized by CoPEC. To support this finding, we examined how CoPEC infection affected tumor growth using MC38 subcutaneous tumor model *in vivo* model. Similarly, tumor volume was negatively correlated with IFNγ, corroborating with data in the right-sided CRC tissues from patients colonized by CoPEC. In support, Lopès et al. ^22^ showed a decrease of CD3^+^ and CD8^+^ T-cells with decreases in the anti-PD-1 immunotherapy efficacy in mice infected by CoPEC. Likewise, a work focused on the spatial effect of the intratumoral microbiota in cancer, reported an increase in CD11b^+^ and CD66b^+^ myeloid cells but lower densities of CD4^+^ and CD8^+^ T-cells in bacteria-positive microniches when compared to bacteria-negative areas. ^5^

Lipid droplet accumulation may modulate signaling pathways that contribute to cell proliferation and chemoresistance^28^. Among different functions, lipid droplets can *i*) to protect membranes from peroxidation reactions under oxidative stress conditions and maintain organelle homeostasis; *ii*) to regulate autophagy by different mechanisms; and *iii*) to respond to exogenous lipid overload to reduce the accumulation of lipotoxic lipids and others^25^. Herein, it is clear that CoPEC infection increases oxidative stress in colon cancer cells which may result in the formation of end-products of ROS-mediated lipid peroxidation. This observation is consistent with macrocyclic Colibactin activity, which induces DNA double-strand breaks via copper-mediated oxidative cleavage. ^56^ Accordingly, we observed that CoPEC colonization heightened the expression of enzymes that are involved in the Land’s cycle when comparing transcripts from the relapsing group. Among those, we found that CoPEC is able to modulate the expression of LPCAT2 enzymes that participate, at different levels, in the regulation of lipid droplet expansion through PC biosynthesis.^52^ LPCAT2 is involved in the reacylation of LPC into PC and is up-regulated by bacterial endotoxin stimulation.^43,44^ Furthermore, we found a lowered infiltration of IFN-gamma-producing CD8^+^ T cells within tumoral area that are populated by bacteria. This is in agreement with the lowered CD8+ T-cell infiltration in tumors with greater expression of LPCAT2 patients that have a poorer prognosis.^51^ In line with altered PC metabolism in CRC^42^, we also noticed an increase in DGKG levels in tumors populated by CoPEC. Several members of the DGK family have been implicated in CRC.^40^ In particular, DGKG plays a role in DNA methylation in CRC tumors, suggesting it is an early event during CRC tumorigenesis. ^41^ Equally of importance, we noticed a change in the expression of enzymes from the cytosolic phospholipase A2 (cPLA2) family involved in the reacylation of the Lands cycle. On the one hand, PLA2G16, PLA2G4D and PLA2G2F were positively regulated as LPCAT2. This result is in agreement with previously published data where cPLA2α is elevated in senescent T cells in the tumor microenvironment.^45^ On the other hand, PLA2G6 (cPLA2 group VI) gene which encodes for a calcium-independent PLA2, was down-regulated in the tumors colonized by CoPEC, accompanied by overexpression of genes mRNAs involved in the decrease of ceramide (e.g. ACER2 and SGMS1). This accords with the lowered generation of ceramide upon inhibition of cPLA2 that enhances colon tumorigenesis.^26^ Cancer cells can use various strategies in response to bacterial genotoxins and probably, a disorder of cPLA2 (increase or decrease) may shift the balance in the colon toward cell growth or survival. In addition, LPC and ceramides were down-regulated in the same conditions. Ceramide is a central molecule of sphingolipid metabolism and is important to maintain cell homeostasis through an orchestrated balance between cell proliferation and apoptosis ^53^. Studies suggest that cancer cells may acquire the ability to maintain ceramide low levels, bypassing normal metabolic control and then, contributing to chemoresistance ^54,55^. These data suggest lipid metabolic reprogramming of tumor cells as a possible alternative route supporting triacylglycerol, potentially through PC and cPLA2 of the Lands cycle. Together, these data suggest that oxidative stress potentially induces a cascade of events in the DNA damage and can trigger lipid droplet accumulation, supporting chemoresistance. It is important to highlight that chemotherapy efficacy is in part a result of its ability to enhance adaptive antitumor immune responses.^57^ Therefore, our results using spatially-resolved metabolomic and transcriptomic approaches clarify how the presence of Colibactin-producing bacteria may locally establish tumor heterogeneity for evading immune surveillance. These findings provide unique insights both for therapeutic intervention and enabling basic research into the mechanisms of CoPEC-induced tumor recurrence.

## MATERIALS AND METHODS

### Patients’ samples

All patients included in the analysis were diagnosed as sporadic cases and with tumors that arise in the right-sided colon. We used 76 samples of the Biobanks that have been set up at the Hospital Henri Mondor (n=29) and at the Hospital of Clermont-Ferrand (n=47). A part of the patients was enrolled in several prospective cohorts named CCR1-3 (Acronyms Valihybritest and Vatnimad; for description, see Sobhani et al. ^58^ and on ClinicalTrials.gov: NCT01270360). This protocol has been approved by the ethics committee of *Comité de Protection des Personnes Paris Est-Henri Mondor* (no. 10-006 in 2010). Another part of these patients underwent surgery for CRC in the Digestive and Hepatobiliary Surgery Department of the University Hospital of Clermont–Ferrand. All patients were adult volunteers and signed informed consent before they were included in the study.

### Identification of CoPEC-positive and negative tumors from RCC patients

We used two methods to validate the presence of *pks* island in the RCC tumor tissue. First, for patients from Hospital of Clermont-Ferrand and Créteil, the samples were analyzed by PCR using specific primers located in the *clbB* and *clbN* genes of the *pks* island: *clbBr* (r for reverse orientation) (5′-CCA TTT CCC GTT TGA GCA CAC-3′), *clbBf* (f for forward orientation) (5′-GAT TTG GAT ACT GGC GAT AAC CG-3′), *clbNr* (5′-CAG TTC GGG TAT GTG TGG AAG G-3′), and *clbNf* (5′-GTT TTG CTC GCC AGA TAG TCA TTC-3′) ^59^.

Posteriorly, for 65 patients from this study, DNA extraction was performed from eight 50 μm cryosections of nitrogen frozen tissue using the QIAamp PowerFecal DNA Kit® (Qiagen, 12830-50) following the manufacturer’s instructions with the following modification: 0.1 mm diameter silica beads were added to the lysis solution provided and shaking was performed at maximum speed for 10 minutes in a vibratory shaker. DNA concentration was measured using the Qubit dsDNA Broad Range assay kit (Invitrogen Q32853). qPCR reactions were performed on a QuantStudioTM 7 Flex Real-Time PCR System (Applied Biosystems 4485701) in 384-well plates with 32ng of DNA, in a final volume of 8µl. Three technical replicates were performed for each sample. Primers and probes listed in the Table were used at a final concentration of 250 nM.

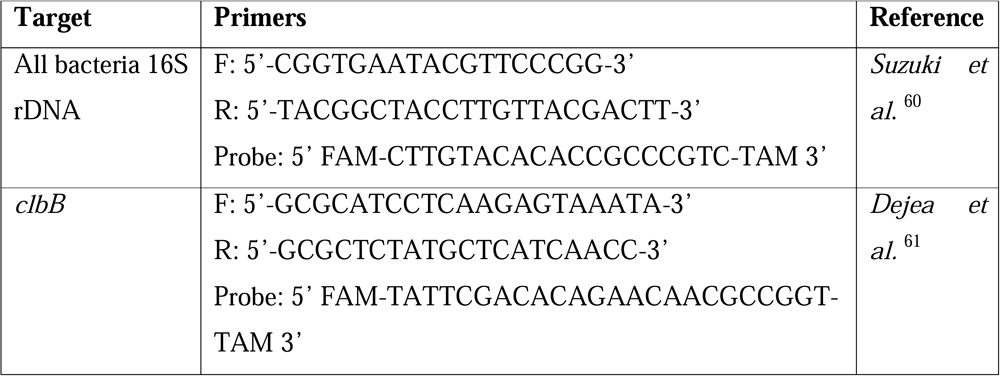

The Master Mix Taqman (Applied Biosystems 4440038) was used with the following amplification steps: 50°C during 2min, 95°C during 15sec with 48 cycles, 60°C during 1min. The amount of *clbB* was determined using the QuantStudioTM Real-Time PCR software (version 1.7.2, Applied Biosystems) and the 2-ΔCt method, with all bacteria 16S rDNA as a calibrator. Patients with amplification were assigned as positive CoPEC.

### Microbiota analysis

Total DNA from 76 tumor tissues was used to perform microbiome analysis. Bacterial 16S rRNA gene V1-V2 region was amplified using 27F and 338R PCR primers and sequenced on a MiSeq (2×250 bp; Illumina, Hayward, CA). Bioinformatic processing and statistical analysis were performed in R software environment as described previously ^62^. Briefly, paired-end fastq files without barcodes and adapters have been quality checked, denoised and prepared for further analysis using the dada2 package ^63^. Bacterial sequencies from amplicon sequence variant (ASV) table were annotated using latest silva database (version 138.1) ^64^. Data normalization and beta-diversity analysis were performed on each taxonomical level using DESeq2 package ^65^. To improve the power of detecting differentially abundant taxa, only those bacteria that appeared in 25% of samples at least in one of the compared groups were used for differential abundance analysis. Differences were considered significant when the corrected p-value (p adjusted) was < 0.05.

### Tissue processing and FISH

Samples of RCC tumor tissue were collected within 30 minutes after surgical resection and immediately frozen in liquid nitrogen and stored at -80°C until further use. This analysis was conducted on 5 μm sections of 12 sporadic RCC patients’ samples using cryostat at -21°C. Serial adjacent sections were obtained for FISH and hematoxylin and eosin (H&E) staining. Tissues were visualized by a pathologist to obtain tumor area and bacteria were stained by FISH using the general bacterial rRNA probe, EUB338 conjugated with Alexa 555 as described previously ^66^.

### Spatial metabolomic using MALDI-FTICR imaging

This analysis was conducted on 10 μm sections of the RCC tissue obtained in a cryostat at -21°C and mounted on ITO slides. On each slide, a quality control section (rat kidney homogenate spiked with Rutine) was added. Imaging of the tumors was performed at 80 μm using a 7T-MALDI-FTICR in full scan mode (75 – 1000) and in negative ion mode. Following the acquisition, the MALDI matrix was removed in a bath of methanol and an H&E staining was performed. In particular, we conducted an *in situ* metabolites analysis dependent on the presence of bacteria in 12 sporadic right-sided CRC patients, who were previously grouped as CoPEC-positive and -negative patients. In addition, quantification was also performed in the total tissue of 65 right-sided CRC patients, acquiring the metabolite intensity of all tumor tissue. The normalized root-mean-square was applied and differential metabolite expression was determined using the limma package (v3.46.0) in R software. The following criteria were applied: p-value produced by Mann−Whitney−Wilcoxon test after FDR correction < 0.05, fold change > 1 or < -1). Volcano plot was used to present differential expressed metabolites with the EnhancedVolcano R package (version 4.0.3). Data acquisition, processing, and data visualization were performed using FlexImaging 4.1 and Multimaging 1.1 (ImaBiotech SAS, France). MSI data were acquired from each tissue section as well as matrix control areas adjacent to the tissue sections to check for analyte dispersion during sample preparation. Metabolite annotation was performed according to Lipid maps (http://www.lipidmaps.org/) and Metlin (https://metlin.scripps.edu).

### RNA sequencing (RNAseq) and CMS classifier from right-sided CRC tumors

Total RNA was extracted from 47 tumor samples using TRIzol®-chloroform extraction method. RNA was sent to the NOVOgene company (China) that carried out the quality control, library preparation, and sequencing. The reads quality was assessed using FastQC (v0.11.4) ^67^ combined with MultiQC (v1.6) ^68^. We performed splice-aware alignment on RNA-seq using the STAR transcriptome aligner (v2.5.0) ^69^ with human genome version GRCh38 from Ensembl release 99 ^70^. After alignment, featureCounts (subread v 1.6.1) was used to obtain the matrix of counts of fragments by genes from sorted BAM files ^71^. We removed samples for which less than 50 % of the fragments mapped to genes and used blast+ (2.9.0) to align the reads on the nt collection and confirm that the lack of alignment to human genes was due to an excess of reads mapping to non-human organisms, suggesting contamination. We filtered gene-level raw counts by removing all genes with no fragment mapped in more than half of the samples. CMS classifications were determined using the CMScaller R package ^72,73^. We used the recommended option ’RNA-seq=TRUE’, which makes CMScaller perform a log2 transformation and quantile normalization and the option ’FDR=0.05’. Normalized counts for further use were generated by applying a quantile normalization using the voom function of the limma package (v3.46.0) ^74^ and Volcano plot was used to present differential expressed genes with the EnhancedVolcano R package (version 4.0.3).

### Cell culture, bacterial infection, lipid droplets, ROS and qRT-PCR

Human HCT116 cells or mouse MC38 cell line, derived from methylcholanthrene-induced C57BL6 murine colon adenocarcinoma cells were used in this study. HCT116 cells were maintained in advanced DMEM F12 and MC38 were maintained in DMEM GlutaMAX. These cells were supplemented with 10% FBS, 1mM L-Glutamine (except to MC38), and Penicillin (100[µ/ml) – Streptomycin (0.1 mg/ml), at 37 °C under 5% CO2 pressure. Cells were routinely tested for mycoplasma. All media components were obtained from Sigma-Aldrich. Treatments were carried out using the clinical CoPEC 11G5 strain isolated from a patient with CRC and its isogenic mutant 11G5Δ*clbQ*, depleted for the *clbQ* gene in the *pks* island and unable to produce Colibactin ^25^. These strains were grown at 37 °C in Luria-Bertani medium overnight. For bacterial infections, cells were infected at a multiplicity of infection (MOI) of 10 or 100 bacteria for 3 h and cells remained in culture for 1h or 5 days after infection. Cells were washed three times with PBS and culture medium with 200 μg/mL gentamicin was added ^17^. Lipid droplets were analyzed using Nile red (Molecular Probes) at 1:2000 in PBS for 15 min and then fixed in 4% paraformaldehyde for 10 min at 4 °C. The slides were mounted in ProLong Gold Antifade with DAPI (Molecular Probes) before imaging. Lipid droplets quantification was performed by counting red lipid bodies on merged pictures (300 cells per cell line). To measure cellular ROS, cells were stained using 5 μM CellRox Deep Red Reagent (Invitrogen, C10422) for 30 min at 37◦C, washed three times with PBS and fixed using 4% paraformaldehyde for 10 min before imaging. The slides were mounted in ProLong Gold Antifade with DAPI (Molecular Probes) before imaging. A semi-quantitative assessment of the percentage of ROS in 300 cells. The scores were as follows: score 0 (absence of CellRox fluorescence); score 1 (weak CellRox fluorescence); score 2 (moderate CellRox fluorescence) and score 3 (strong CellRox fluorescence). Data are expressed by the percentage of cells in each score and group (two independent experiments and duplicates for each experiment). Images were acquired with an Axio Imager M2 (Zeiss) coupled with an Apotome.2 (×63 or x100 objective).

### SpiderMass analysis

After almost 70% confluence, HCT116 and MC38 cells were washed two times with DPBS, dried under PSM for 10 min at room temperature then analyzed by the SpiderMass directly into the cell plate. The overall layout of the SpiderMass setup has already been covered elsewhere ^75^. In brief, the system is made up of three parts: the mass spectrometer itself, a laser system for remote micro-sampling of tissues and a transfer line allowing for the transfer of the micro-sampled material. In this study, the laser intensity was set to 4 mJ/pulse and a 200 µL/min infusion of isopropanol was administered during each acquisition. 200 µg/mL of Leucine enkephalin was added to the infusion to play the role of a lockmass. The acquisition was composed of a burst of 10 laser shots resulting in an individual spectrum. Spectral acquisition was performed both in negative ion mode in sensitivity mode and the mass range was set to m/z 50-2000. The raw files were imported into “Abstract Model Builder” - AMX (version 1.0 1972.0, Waters, Hungary) to perform multivariate statistical analyses using linear discriminative analyses (LDA). Discriminative ions were found looking at each dual condition loading plot. Boxplots for each specific ions were obtained thanks to Kruskal-Wallis significant tests.

### RNAseq analysis of MC38 cell line

MC38 cells were infected using the 11G5 strain and 11G5Δ*clbQ* strain at a MOI of 100 bacteria during 3 h and cells remained in culture for 1h after infection. Non-infected cells also were analyzed under the same conditions. Posteriorly, cells were immediately lysed in RLT buffer and RNA was extracted using RNeasy Mini Kit (QIAGEN) following manufacturer protocol. mRNA library preparation was realized following the manufacturer’s recommendations (Illumina Stranded mRNA Prep Kit from ILLUMINA). Final samples pooled library prep was sequenced on Novaseq 6000 ILLUMINA with S1-200cycles cartridge (2x1600Millions of 100 bases reads) corresponding to 2x30Millions of reads per sample after demultiplexing.

### Cell viability

Cell viability was investigated using WST-1 assay reagent (ab155902). This test is based on the cleavage of the tetrazolium salt WST-1 to formazan by cellular mitochondrial dehydrogenases. 5×10^3^ HCT116 cells were infected with 11G5 strain and 11G5ΔclbQ strain at MOI of 500 bacteria during 3 h and cells remained in culture for 7 days after infection. Non-infected cells were used as a negative control. All cells were treated with 0 and 5 μg/mL of oxaliplatin in 96-well plate at 37°C for 72h. WST-1 solution was added and incubated for 4h at 37°C. Absorbance was measured using FLUOstar Omega plate reader (BMG Labtech) at 420 nm. The percentage of cell viability was calculated in relation to non-infected cells without oxaliplatin.

### Spatial transcriptomic using RNAscope *in situ* hybridization

RNAscope multiplex fluorescent reagent 2.5 HD kit assay (Advanced cell diagnostics, Newark, CA, USA) was performed using 10 frozen RCC tissue (5 samples for each CoPEC-positive and negative group) and cryostat cut sections of 10 µm were collected and mounted onto Superfrost™ Plus microscope slides. Slides were immersed in the 4% PFA for 1h at 4°C. Posteriorly, the sections were dehydrated and slides were baked for 1h at 60°C and a hydrophobic barrier was drawn around the tumor tissue. Endogenous peroxidases were blocked using a hydrogen peroxide solution for 10 min at room temperature and then, was used RNAscope proteas IV sob under the same conditions. The probes hybridization process was performed using the HybEZ™ II oven for 2h at 40°C for Hs-IFNγ (310501, Opal 570) and Hs-CD8A (560391-C3, Opal 520) probes. Between each amplification and staining step, slides were washed twice in 1X RNAscope wash buffer for two minutes. Then, the slides were incubated with DAPI (ThermoFisher Scientific) for 3 min at room temperature and placed with ProLong Gold antifade mounting solution (ThermoFisher Scientific) prior to imaging. Programmed cell counting using Imaris (Bitplane version 9.5.0) in order to count the number of spots for CD8A (5000 – 7000 cells, approximately) and for CD8A and IFNγ in the same cell (500 - 1000 cells, approximately).

### Mice infection and tumor growth

Animal protocols were approved by the Ministère de l’Education Nationale, de l’Enseignement Supérieur et de la Recherche (APAFIS#20990). This study was performed using male wild-type C57BL/6J mice. All mice were housed in conventional conditions at the animal care facility of the Institute Pasteur of Lille and had unlimited access to food and water. The colonization by CoPEC using wild-type (WT) mice was conducted as described previously^76^. In brief, to enhance *Escherichia coli* strain colonization, we administered streptomycin (2.5 g/l) for 3 days prior to oral inoculation with 11G5 strain (n=7) or its isogenic mutant 11G5Δ*clbQ* (n=6) (≈ 1 × 10^9^ bacteria in PBS). Eight days after infection, to induce tumor formation, 5×10^5^ MC38 cells in PBS were injected subcutaneously into the left flank of male mice. Tumor volume in mm^3^ was monitored two or three times a week by the measurement of two perpendicular diameters using a caliper according to the formula L×S2/2, where L and S are the largest and smallest diameters in mm, respectively.

### RNA extraction and qRT-PCR from mice samples

Tumor tissues were homogenized with ceramic beads on a MagNA Lyser (Roche). RNA was extracted using RNeasy Mini Kit (QIAGEN) following manufacturer protocol. 250ng RNA of each sample was retro-transcribed using Affinity Script cDNA synthesis kit (Agilent Technologies). 5ng of cDNA was used for the qRT-PCR reaction using Brilliant III Ultrafast SYBR Green QPCR master mix (Agilent Technologies) on AriaMx qRT-PCR system (Agilent Technologies). The following primers were used: mIFN-γ For= GCTTTGCAGCTCTTCCTCAT and Rev= CCAGTTCCTCCAGATATCCAAG; mLpcat2 For=TCCCAGAAGGTACTTGTACTAATCG and Rev=TGTTTGGGTATCTGAGGAGGA.

### Statistical Analysis

Progression-free survival (PFS) was evaluated using Kaplan–Meier method available in the R (version 4.0.3) package survival. The p values are from log-rank tests. Pearson correlation was used to investigate the correlation between PC and CD8^+^ T-cell spots with R software (version 4.0.3) and GraphPad Prism software (version 6.0). Statistical analyses between two groups were performed with the student’s t-test or a Mann-Whitney U test, conforming to the results of the normality test. A one-way ANOVA followed by a post-test Bonferroni correction also was used when appropriate, using GraphPad Prism software (version 6.0). A p value less than 0.05 was considered statistically significant.

## Acknowledgments

We would like to thank the BioImaging Center Lille platform for assistance with image acquisition. We thank monitoring platform studies of URC St Antoine (Pr T Simon for Vatnimad PHRC study), URC Henri Mondor (Pr F Canoui-Poitrine; for Valihybritest ANR study), Biomics Platform, C2RT, Institut Pasteur, Paris, France, supported by France Génomique (ANR-10-INBS-09), Institut pour le Recherche sur le Cancer de Lille (IRCL), I-Site ULNE, INSERM (Messidore) and IBISA. This work benefited from equipment and services from the iGenSeq core facility (Genotyping and sequencing), at ICM.

## Contributors

NOA and MC performed study design, acquisition of data, interpretation of data and statistical analysis. OB, SA, LA and NOA performed bacterial cultivation, cell culture and qRT-PCR analysis. AV, IN, CM, TP and MM acquired, analyzed and interpreted data and statistical analysis from RNAseq. LM, DN and SK analyzed, interpreted data and statistical analysis of microbiota data. RR performed FISH test. EB,DL,EL and AM analyzed and interpreted data from qPCR to identification of colibactin. LL, IF and MS conducted SpiderMass analysis. GD, PS, JG, DP, CG, NB and RB acquired, analyzed and interpreted data from Clermont-Ferrand patients. DM and IS acquired, analyzed and interpreted data from Créteil. NOA wrote the manuscript and all authors discussed the results and commented on the manuscript. MC supervised the entire project.

## Ethics approval

All animal experiments were approved by the local investigational review board (APAFIS#20990). Animal studies were performed in an accredited establishment (N° B59-350009) according to governmental guidelines N°86/609/CEE.

## Data available statement

All data relevant to the study are included in the article or available as online supplemental information. The data not shown will be shared on reasonable request to the corresponding authors.

## Supplementary

**Figure S1. CoPEC colonization lowered the bacterial richness in the ASV-based dataset.** Alfa-diversity analysis results between positive and negative patients colonized by colibactin-producing *Escherichia coli* (CoPEC) on (A) observed ASV index, (B) Fisher’s alpha diversity index, (C) Chao1 estimated richness, and (D) Pielou evenness.

**Figure S2. Beta-diversity analysis results of CMS3 group between positive and negative patients colonized by colibactin-producing *Escherichia coli* (CoPEC).** The left plot represents the log_2_ fold change value of the mean normalized abundance ratio of positive and negative CoPEC groups; the middle plot represents the p-value obtained by deseq2 differential abundance analysis; the right plot represents the mean normalized abundance for each significant abundance bacteria.

**Figure S3. FUMA results of the overrepresented genes in response to Colibactin.** Significantly enriched gene sets from the Gene Ontology (GO) Biological Process category of the 128 up- (C) and 126 down- (D) regulated genes expression for RCC patients colonized by CoPEC. The blue bars represent the enrichment p-value (-log10 adjusted) after FDR correction considering the number of gene sets tested per category. The red bars indicate the proportion of overlapping inputted genes according to the size (number of genes) of each gene set. The orange squares show the inputted genes that are part of the enriched gene sets

