## Supplementary figures and images for "The Colibactin-Producing *Escherichia coli* alters the tumor microenvironment to immunosuppressive lipid overload facilitating colorectal cancer progression and chemoresistance"

### Supplementary Figures

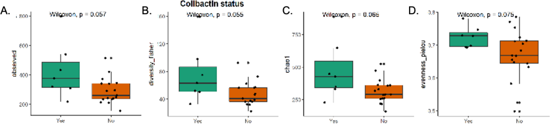

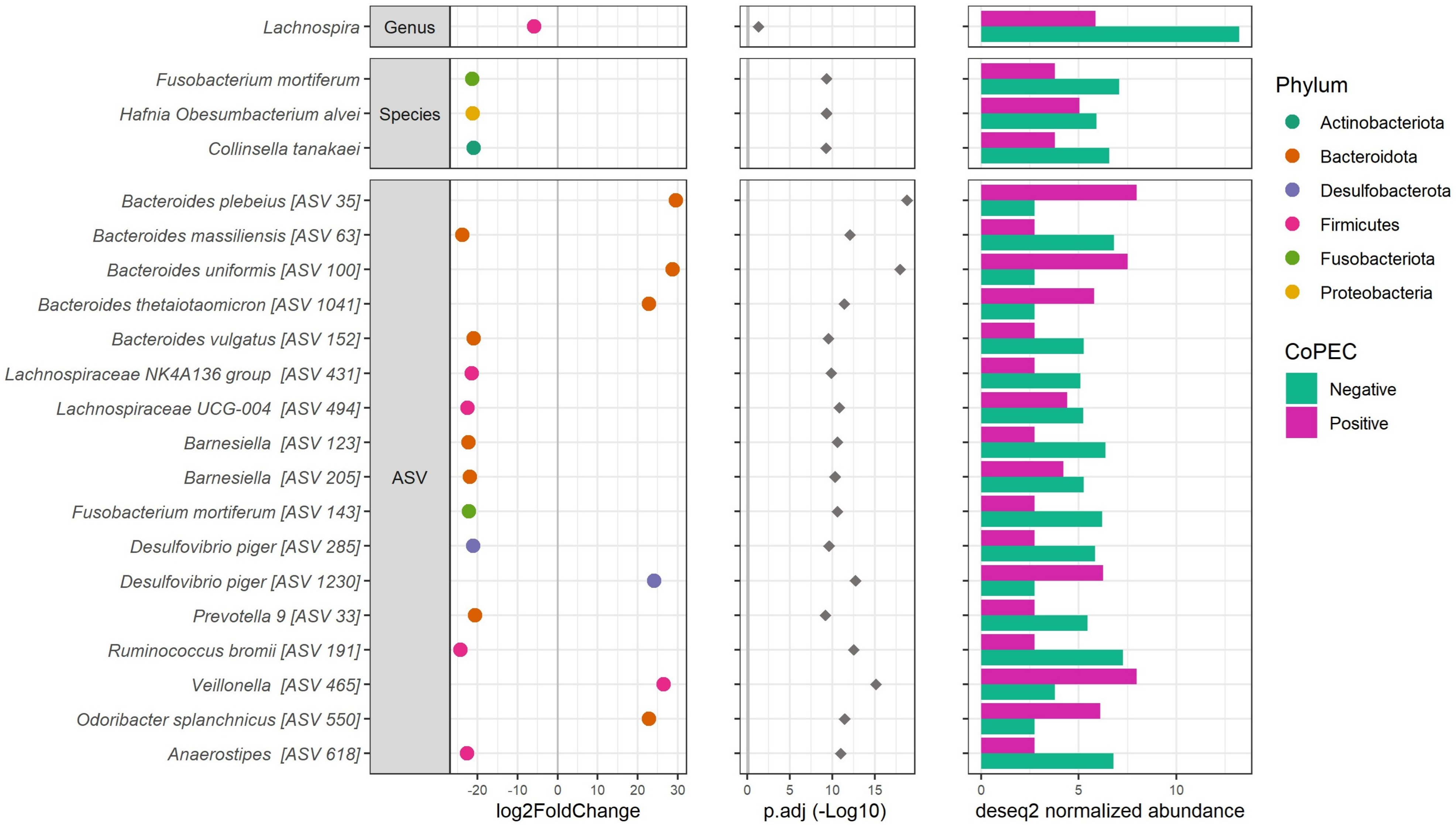

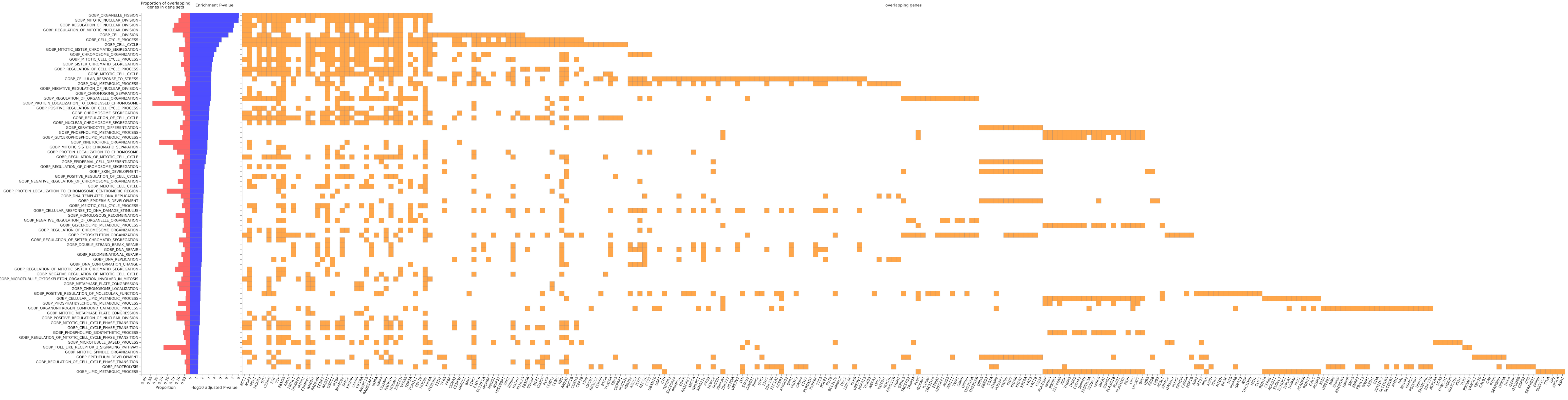
