## Supplementary Table for "The Colibactin-Producing *Escherichia coli* alters the tumor microenvironment to immunosuppressive lipid overload facilitating colorectal cancer progression and chemoresistance"

**Table S1: List with discriminant ions (400 – 1100 m/z) corresponding to lipids from three conditions: 11G5-, 11G5∆*clbQ*-infected HCT116 and non-infected cells.**

| ***m/z*** | **11G5** | **11G5ΔclbQ** | **Non infected** | **Potential lipids** |
| --- | --- | --- | --- | --- |
| 542.45 | **-** | **+** |  | Cer 32:0;O4 |
| 546.45 | **-** | **+** |  | LPC 20:2 |
| 560.45 | **-** | **+** |  | CerP 30:1;O2 ou Cer 36:3;O2 |
| 570.45 | **-** | **+** |  | LPC 22:4 |
| 574.45 | **-** | **+** |  | LPC 22:2 |
| 594.55 | **+** | **-** |  | Cer 38:0;O2 |
| 603.45 |  |  | **+** | [DG (36:10)-H]^-^ |
| 666.45 |  | **+** | **-** | [PE (O-32:5)-H]^-^ |
| 682.55 | **-** | **+** |  | [Cer (t42:0(2OH))-H]^-^ |
| 684.65 |  | **+** | **-** | Not identify |
| 694.45 | **+** |  | **-** | [PE (O-34:5)-H]^-^ |
| 740.55 | **-** |  | **+** | [PE (36:3)-H]^-^ |
| 742.55 | **-** |  | **+** | [PE (36:2)-H]^-^ |
| 800.45 | **+** | **-** |  | [PS (38:9)-H]^-^ |
| 802.55 | **+** |  | **-** | [PC (38:7)-H]^-^ |
| 822.55 | **+** | **-** |  | [PS (O-40:5)-H]^-^ |
| 824.55 | **+** | **-** |  | [PS (O-40:4)-H]^-^ |
| 884.45 | **+** | **-** |  | [PS (44:9)-H]^-^ |
| 888.65 |  | **+** |  | [PS (44:7)-H]^-^ |
| 910.55 | **-** | **+** |  | [PS (42:0)-H]^-^ |
| 913.45 | **+** |  | **-** | [PI (40:4)-H]^-^ |

+ (up-regulated); - (down-regulated)

**Table S2: List with discriminant ions (600 – 1100 m/z) corresponding to lipids from three conditions 11G5-, 11G5∆clbQ-infected MC38 and non-infected cells.**

| ***m/z*** | **11G5** | **11G5ΔclbQ** | **Non infected** | **Potential lipids** |
| --- | --- | --- | --- | --- |
| 680.55 | **-** |  | **+** | [Cer (t42:2(2OH))-H]- |
| 682.55 | **-** | **-** | **+** | [Cer (t42:0(2OH))-H]- |
| 768.55 | **-** |  | **+** | [PE (38:3)-H]- |
| 844.65 | **-** |  | **+** | [PE (44:7)-H]- |

+ (up-regulated); - (down-regulated)
